# Genome-wide selection scans integrated with association mapping reveal mechanisms of physiological adaptation across a salinity gradient in killifish

**DOI:** 10.1101/254854

**Authors:** Reid S. Brennan, Timothy M. Healy, Heather J. Bryant, Man Van La, Patricia M. Schulte, Andrew Whitehead

## Abstract

Adaptive divergence between marine and freshwater environments is important in generating phyletic diversity within fishes, but the genetic basis of adaptation to freshwater habitats remains poorly understood. Available approaches to detect adaptive loci include genome scans for selection, but these can be difficult to interpret because of incomplete knowledge of the connection between genotype and phenotype. In contrast, genome wide association studies (GWAS) are powerful tools for linking genotype to phenotype, but offer limited insight into the evolutionary forces shaping variation. Here, we combine GWAS and selection scans to identify loci important in the adaptation of complex physiological traits to freshwater environments. We focused on freshwater (FW)-native and brackish water (BW)-native populations of the Atlantic killifish (*Fundulus heteroclitus*) as well as a population that is a natural admixture of these two populations. We measured phenotypes for multiple physiological traits that differ between populations and that may contribute to adaptation across osmotic niches (salinity tolerance, hypoxia tolerance, metabolic rate, and body shape) and used a reduced representation approach for genome-wide genotyping. Our results show patterns of population divergence in physiological capabilities that are consistent with local adaptation. Selection scans between BW-native and FW-native populations identified genomic regions that presumably affect fitness between BW and FW environments, while GWAS revealed loci that contribute to variation for each physiological trait. There was substantial overlap in the genomic regions putatively under selection and loci associated with the measured physiological traits, suggesting that these phenotypes are important for adaptive divergence between BW and FW environments. Our analysis also implicates candidate genes likely involved in physiological capabilities, some of which validate *a priori* hypotheses. Together, these data provide insight into the mechanisms that enable diversification of fishes across osmotic boundaries.

**Author Summary:** Identifying the genes that underlie adaptation is important for understanding the evolutionary process, but this is technically challenging. We bring multiple lines of evidence to bear for identifying genes that underlie adaptive divergence. Specifically, we integrate genotype-phenotype association mapping with genome-wide scans for signatures of natural selection to reveal genes that underlie phenotypic variation and that are adaptive in populations of killifish that are diverging between marine and freshwater environments. Because adaptation is likely manifest in multiple physiological traits, we focus on hypoxia tolerance, salinity tolerance, and metabolic rate; traits that are divergent between marine and freshwater populations. We show that each of these phenotypes is evolving by natural selection between environments; genetic variants that contribute to variation in these physiological traits tend to be evolving by natural selection between marine and freshwater populations. Furthermore, one of our top candidate genes provides a mechanistic explanation for previous hypotheses that suggest the adaptive importance of cellular tight junctions. Together, these data demonstrate a powerful approach to identify genes involved in adaptation and help to reveal the mechanisms enabling transitions of fishes across osmotic boundaries.

## Introduction

Evolutionary transitions from marine to freshwater environments have served a vital role in generating the phyletic diversity within ray-finned fishes [1,2]. It has been hypothesized that adaptive divergence of estuarine species in different salinities is a major driver of these transitions [3]. Indeed, adaptive divergence is a major component in the generation of new species across diverse phyla (ecological speciation [4]). Therefore, discovering the mechanisms that enable such adaptive divergence between osmotic niches is important for understanding the mechanisms that drive speciation within the fishes. To understand the processes involved in this divergence, it is necessary to understand how traits and their underlying molecular pathways and genes facilitate adaptation and isolation in different ecological niches.

Despite its importance, identifying the mechanistic basis of adaptation is often difficult. Many recent successes include morphological traits, for example in stickleback fish, field mice, and butterflies [7-9], and these studies have implicated a small number of genes of major effect in these processes. However, it has been argued that these classic examples may not be typical of how evolution often proceeds [10]. For many adaptive challenges, relevant traits may be highly polygenic, and multiple traits may contribute to a complex adaptive phenotype. This may be especially true along marine to freshwater gradients where many environmental features vary (e.g., salinity, oxygen, pH, food, predators, pathogens, symbionts, etc. [11-14]) and may affect fitness. Genome wide association mapping (GWAS) is often deployed to identify genetic variation underlying phenotypic traits, but is often inefficient for discovering variation associated with highly polygenic traits [10]. Also, GWAS offers limited insight into the fitness effects of that variation. In contrast, genome-wide scans for signatures of selection offer an unbiased approach for discovering the many genomic regions that underlie local adaptation. However, selection scans rarely enable inference that links specific genotypic variation with relevant phenotypes. In confronting this conundrum, recent work has suggested integrating GWAS and genome-wide scans for signatures of natural selection [15]. By combining the approaches, one can amplify the polygenic signal from GWAS, and identify causal variants for specific phenotypes while simultaneously addressing whether or not these variants are the targets of selection [16]. Since adaptation to fresh water should require the modulation of multiple traits, fishes are an ideal system for understanding how complex physiologies evolve to enable divergence across key ecological boundaries.

Killifish within the *Fundulus* genus provide a particularly compelling model system for studying diversification and adaptation between marine and freshwater environments. Within this genus, marine to freshwater transitions have occurred at least three times independently [17,18]; such lability is rare for closely related species [2]. *Fundulus heteroclitus* is a particularly good microevolutionary model species to identify the genetic and physiological mechanisms important for freshwater adaptation. This species inhabits environments ranging from marine to fresh water [19], and exhibits adaptive divergence along salinity gradients [20]. In the Potomac River and Chesapeake Bay, genetically distinct populations occupy different osmotic niches. A genetic break occurs along a salinity cline where individuals upstream of the freshwater boundary are distinct from downstream fish in brackish water [20,21]. Freshwater populations (FW-native) exhibit environmentally-dependent physiological differences from their brackish water counterparts (BW-native) when acclimated to freshwater, potentially representing the incipient stages of speciation across an osmotic boundary [20,22,23]. This is an ideal system to identify genomic regions under divergent selection. Because these populations are recently diverged, genome-wide association mapping can be used to identify loci that are likely to influence physiological performance. Integrated together, these approaches can reveal the genes important for adaptation and divergence in freshwater environments.

In this study, we seek to identify the physiological and genomic basis of adaptation to fresh water. Specifically, we addressed the following questions: 1) Are adaptive physiological traits divergent between BW-native and FW-native populations?; 2) What is the genetic basis of freshwater adaptation? To address these questions, we sampled *F. heteroclitus* individuals from BW-native and FW-native populations (“parental” populations) along a salinity gradient in Chesapeake Bay, and individuals that are genetically intermediate to BW‐ and FW-native fish (“admixed” population; Fig. 1). The presence of admixed individuals can be leveraged to identify genomic regions associated with phenotype (admixture mapping [24]). Furthermore, we can test for the independent segregation of phenotypic traits in admixed individuals to understand the modularity of traits and how this may influence adaptation. To quantify physiological divergence between populations, all individuals were phenotyped for hypoxia tolerance, metabolic rate (MR), and salinity tolerance – traits known to differ between populations of this species [23,25,26]. We also compared morphology since that has been shown to adaptively diverge between marine and freshwater habitats in other fish model systems [27]. We genotyped these individuals using RADseq and performed selection scans to identify genomic regions evolving by natural selection. We performed association mapping to identify the genetic variants underlying variation in physiological traits. This combination of selection scans and association mapping is especially powerful. Regions overlapping between the approaches are those that are mostly likely to be involved in adaptive divergence of this suite of physiological traits between marine and freshwater environments.

**Figure 1.**
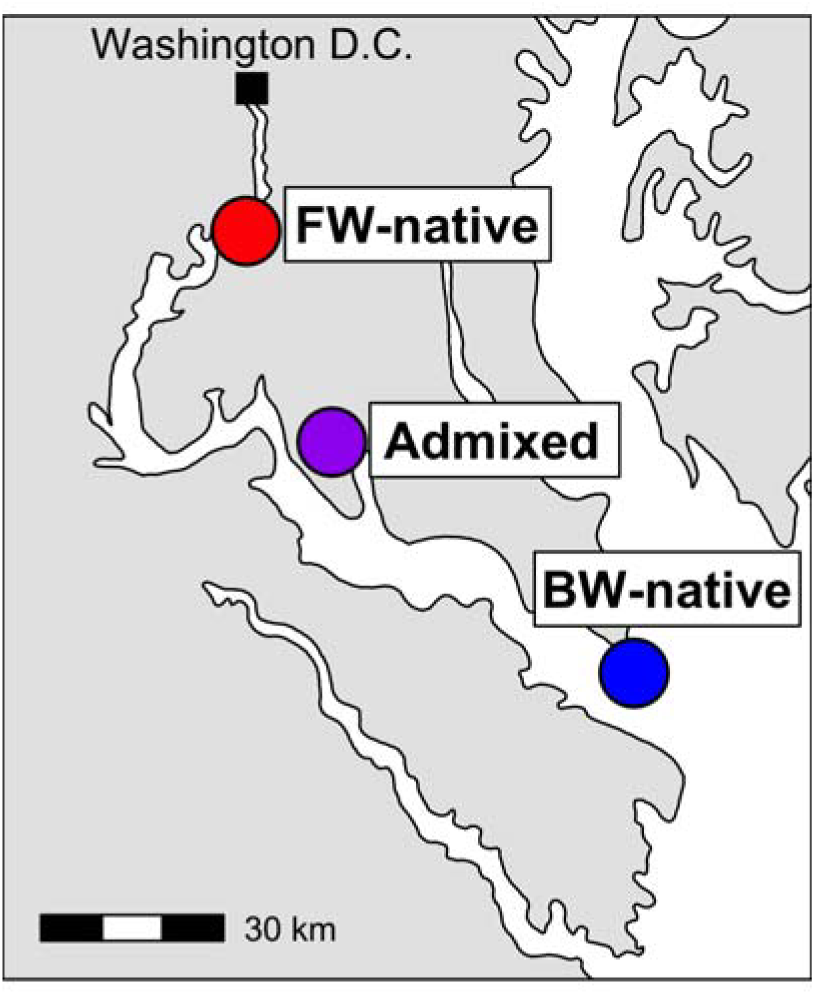
Map of sampling locations in the Potomac River and Chesapeake Bay.

## Methods

### Fish collection and lab acclimation

Adult killifish were collected from the FW-native (n = 40), BW-native (n = 40), and admixed (n = 257) populations in June 2014 (Fig. 3.1). FW-native individuals were sampled from Piscataway Park, near Accokeek, MD (38°41′42.18″N, 77°3′10.38″W) and BW-native fish were from Point Lookout State Park, near Scotland, MD (38°3′10.90″N, 76°19′34.38″W). The admixed population was collected at Allen’s Fresh Run, MD (38°21′54.54;″N, 76°58′52.02″W). Water parameters have been measured by the Chesapeake Bay Program since 1984 and are freely available at http://data.chesapeakebay.net/. The sampling stations closest to our collection sites were used to calculate dissolved oxygen and salinity for each population. The corresponding sampling locations names for BW-native, admixed, and FW-native populations are LE2.3, RET2.4, and TF2.1, respectively. Mean dissolved oxygen levels over the past 30+ years for each site are as follows (in mg/l O_2_ ± standard deviation): BW-native 7.14 ± 3.75, Admixed 7.19 ± 2.97, FW-native 8.69 ± 2.53. Salinity was calculated from the same locations (in parts per thousand(ppt) ± standard deviation): BW-native 14.86 ± 3.15, Admixed 7.80 ± 3.42, FW-native 0.00 ± 0.01. Additionally, the maximum salinity at the FW-native site over the past 30+ years of monitoring is 0.12 ppt demonstrating that this site is exclusively fresh water.

Fish were shipped to the University of British Columbia, Canada and held in a recirculating aquarium system in 200 L tanks at 15 °C and 12L:12D. Acclimation salinity was 20 ppt, made using dechlorinated City of Vancouver tap water adjusted with Instant Ocean® Sea Salt (Instant Ocean, Spectrum Brands, Blacksburg, VA). Each individual was marked with a unique fluorescent elastomer tag (Northwest Marine Technology, Shaw Island, WA) and held separated by population. Nutrafin® Max Tropical Fish Flakes (Hagen, Mansfield, MA) were used for feeding, though a 24 h fast was implemented prior to any experimental measurements. Following each phenotypic measure, acclimation conditions were re-established for at least one month before the next measure.

### Phenotype data

Three physiological traits were characterized for each fish, including salinity tolerance, hypoxia tolerance, and MR. Routine oxygen consumption was used to as a proxy for MR and measured using closed respirometry [26,28]. Repeatability of individual MR was within 10%, as assessed from preliminary tests. For the FW-native, BW-native, and admixed population, data were collected for 36, 40, and 218 individuals. Hypoxia tolerance was calculated as the amount of time required for an individual to lose equilibrium in 0.2 kPa O_2_ water [25] where sample size was 40, 40, and 248 for FW-native, BW-native, and admixed populations. Salinity tolerance was measured using plasma chloride homeostasis following an acute fresh water exposure. Fish were transferred from 20 ppt to 0 ppt and allowed to acclimate for 24 h. At 24h, individuals were sacrificed by pithing and blood was collected by caudal puncture using heparinized hematocrit capillary tubes. Blood was immediately spun at 1500 g for 1 minute to isolate plasma. Blood plasma was snap frozen in liquid nitrogen for later Cl^-^ quantification. Cl^-^ was used to assess the salinity tolerance as it is the ion that most consistently reveals divergence in regulation between *F. heteroclitus* populations from different osmotic environments; we sampled fish at 24 h post-transfer because it is at this time that population differences are greatest [29,30]. Cl^-^concentrations were quantified using a colorimetric mercuric thiocyanate method [31,32]. Finally, we characterized morphology for 12 landmarks (Fig. S5) of 24 FW-native, 29 BW-native, and 152 admixed individuals using TpsDig [33]. For individual fish, data were collected for all three physiological parameters from 38, 34, and 210 FW-native, BW-native, and admixed individuals, respectively.

### Phenotype statistics

Physiological data were analyzed in an ANOVA framework in R with *population* as a main effect. Mass had an effect on all phenotypes and was included as a covariate in the model. However, residuals from these models failed the assumption of normality, therefore we re-ran all statistics using the non-parametric Kruskal-Wallis rank sum test. Both models showed the same significant results, therefore we present the results from only the parametric analyses. For clarity of figures, we present mass-corrected values from linear models where salinity and hypoxia phenotypes are scaled to a fish size of 4 g. Correlations between phenotypes were assessed in admixed individuals only. Regressions between hypoxia, MR, and salinity tolerance were performed in all possible combinations. Because mass was slightly different for each phenotype (because of growth during the time between physiological measures) we mass corrected phenotypes to a standardized 4 g fish before investigating correlations.

Morphological data were analyzed using MorphoJ [34]. Landmark positions were imported to MorphoJ and a Procrustes fit was performed. The effects of allometry were controlled for by a regression of Procrustes coordinates and centroid size and residuals were obtained for further analyses. Shape variation was summarized with a PCA on residuals and differences between groups were calculated using canonical variates analysis (CVA). Positions from PC1 and PC2 were used for subsequent association mapping.

### Genotyping

A custom DNA extraction method utilizing Agencourt Ampure XP beads (Beckman Coulter) was used to obtain genomic DNA for each individual. Details of the method are provided in Ali et al. [35]. Briefly, fin clips were digested overnight at 56 °C in 4.2 mg/ml Proteinase K followed by an Ampure XP bead cleanup and elution in low TE (10 mM Tris–HCl, pH 7.5, 0.1 mM EDTA). Genotyping was performed using restriction site-associated DNA sequencing (RADseq) following the protocol developed by Ali et al. [35]. For each sample, 250 ng DNA was digested with the restriction enzyme *Sbf*I. Individual barcodes were annealed to each sample by adding 2 μl indexed SbfI biotinylated RAD adapters (50 nM). These adapters feature 96 unique 8-bp barcodes that distinguish individuals within a single library. Libraries of 96 individuals were then pooled and fragmented to 200-500 bp using a Bioruptor NGS sonicator with 9 cycles 30 sec on/90 sec off. We assessed fragment size using a 2% sodium borate gel and further fragmentation was performed as needed. Dynabeads M-280 streptavidin magnetic beads (Life Technologies, 11205D) were used to physically isolate the RAD-tagged DNA fragments from off-target DNA and the final DNA was eluted in 55.5 μl low TE. These “RAD libraries” were then prepared for Illumina sequencing using a NEBNext Ultra DNA Library Prep Kit for Illumina. During this process, each RAD library of 96 individuals receives a unique barcode, which allows multiplexing of RAD libraries during sequencing.

Libraries were sequenced using 150-bp paired end reads on an Illumina HiSeq 4000 at the University of California Davis Genome Center. A single lane was first sequenced to aid with normalization prior to definitive library sequencing. From this preliminary single lane of data, libraries were demultiplexed by Illumina barcode then by RAD barcode. RAD barcode demultiplexing was performed using custom Perl scripts that require a perfect match of the barcode and partial restriction site. Reads that contained barcodes on both the forward and reverse reads were discarded. The number of reads per individual was counted and used to renormalize libraries and resequence. We went back to the original fragmented and barcoded DNA from the RAD preparation (prior to physical isolation of the RAD libraries) and adjusted the amount of DNA taken from each individual. This step ensures equal depth of sequencing across all individuals. This DNA was then re-pooled as above and sequenced on an additional two lanes of an Illumina HiSeq 4000. These data were demultiplexed as described above.

Reads were aligned to the *F. heteroclitus* reference genome 3.0.2 [36] using BWA-MEM version 0.7.12 [37] and PCR duplicates were marked using SAMBLASTER version 0.1.22 [38]. Variants were called using Freebayes (v0.9.21-19-gc003c1) where reads with a mapping quality < 30 and discordantly and duplicate reads were discarded [39]. We used GATK [40] to filter SNPs based on coverage, where each site required 80% of the samples to have at least 8x coverage. VCFtools version 0.1.13 [41] was used to filter the resulting variants to retain only sites with bi-allelic single nucleotide polymorphisms (SNPs), base quality >20, minor allele frequency > 0.01, and average coverage per individual less than 100x. Nine individuals had less than 50% of the average read depth and were removed from the analysis. Variants were required to be in Hardy-Weinberg equilibrium in at least 2 of the 3 populations. This resulted in 139,721 high quality SNPs that were used in subsequent analyses. Variants were converted to their appropriate position on the *F. heteroclitus* genetic map (unpublished data). All variants were retained, regardless of their inclusion in the genetic map. Of these, 126,651 (90.6%) were located on assembled chromosomes.

### Population genetics data analysis

To understand the genetic relationships between populations, we took two approaches: principal component analysis (PCA) and admixture analysis. Variants were thinned to remove linkage disequilibrium (LD) based on the variance inflation factor in PLINK v1.90 [42] with the options ‐‐*indep 50 5 2*. PCA was run using PLINK v1.90. Admixture analysis was run with 2 ancestral populations in Admixture v1.23 [43], which calculates the maximum likelihood of the ancestry of each individual.

To identify loci putatively under selection, we used a combined F_ST_ and π outlier approach, which are common approaches to identify regions under selection [44] and indicate elevated allelic differentiation and reduced genetic diversity, respectively. Weir and Cockerham’s F_ST_ was calculated per site using VCFtools v0.1.13. Values for π were calculated using Stacks v1.46 and are similar to expected heterozygosity [45,46]. We sought to identify genomic regions with signatures of selection specifically in the derived FW-native population. Therefore, we calculated π within each population then took the difference between the FW-native and the BW-native population. In this case, more negative values (i.e., lower π in the FW-native population) are evidence of a selective sweep after entering fresh water. F_ST_ and π were combined using a multivariate outlier approach as implemented in the R package MINOTAUR [47]. Raw statistics were converted to rank based p-values based on a uniform distribution, reflecting quantile values from the empirical distribution [48]. To identify selection in fresh water, we based these p-value conversions on right-tailed and left-tailed expectations for F_ST_ and π, respectively. This identifies variants that have high F_ST_ between the populations and lower π in the FW-native individuals. These p-values were then negative log_10_ transformed and the Mahalanobis distances for all SNPs were calculated. Mahalanobis distance represents the number of standard deviations a point lies from the center of a multivariate distribution. Importantly, this measure does not assume independence between statistics, though it does assume smooth dispersal from a centroid, making it necessary to perform the aforementioned p-value transformations. This approach follows the Md-rank-P method described in Lotterhos et al., [48]. From this distribution, we considered the top 1% of SNPs to be outliers, and therefore the most likely candidates as targets of natural selection.

### GWAS

Association mapping was conducted for each phenotype using genome-wide efficient mixed-model association (GEMMA) [49]. We implemented univariate linear mixed models (LMM) while incorporating a n x n relatedness matrix as a random effect. GEMMA was used to generate the centered relatedness matrix using all markers and all individuals. Because different numbers of individuals were phenotyped for each trait (see “Phenotype Data”, above), a small number of SNPs had MAFs that fell below our 0.01 cutoff and were removed from the analysis. As such, for hypoxia tolerance, MR, and salinity tolerance 139 451, 139 242, and 138 951 SNPs, respectively, were included in the LMM. Admixture proportions and log transformed individual mass were included as covariates for single SNP associations and Wald test p-values for log transformed phenotypes were obtained. Admixture proportions were from the admixture analysis described above. Genome-wide significance was calculated based on the number of SNPs that are not in LD (76 655) and Bonferroni correction with α = 0.05.

### Identification of candidate loci

We sought to identify loci that were diverging by natural selection between marine and freshwater environments, and also associated with physiological phenotypes that differed between BW-native and FW-native populations. However, the physiological phenotypes we focused on are complex and likely polygenic, making it difficult to identify genetic markers that are statistically significantly associated with phenotype – a well-known challenge of GWAS [10]. Therefore, we chose a liberal approach to identify candidates by identifying the SNPs falling in the lowest 0.1% of the distribution as GWAS outliers. The goal in this approach is to provide a subset of loci that are most likely involved in the phenotypes investigated. This 0.1% cutoff corresponds to the following p-values: salinity tolerance: 0.001, hypoxia tolerance: 0.0009, MR: 0.0007. We determined the distance at which linkage disequilibrium decayed to background levels by comparing inter-chromosome values (R^2^ = 0.0039) to intra-chromosome values. Decay in LD with physical distance was calculated using non-linear regression [50,51] implemented in the nls package in R [52]. Using this approach, R^2^ values decayed to background within 27 kb. It should be noted that this is likely an upper bound. Using a more conservative measure, the point at which R^2^ drops below 0.1, LD decays after 5 kb [53].

Outlier loci (F_ST_ and π signals integrated by Mahalanobis distance) were merged into outlier windows when any two SNPs within 27kb were top outliers; this value was chosen based on the calculated decay in LD. The overlap between windows showing signatures of selection and GWAS outliers was then identified using Bedtools v2.17.0 [54]. We calculated the probability of obtaining the observed overlap by chance using a permutation test. Minor allele frequency (MAF) may influence the probability that an allele is an outlier in the selection scan and may also influence which GWAS variants overlap with outlier regions. Allele frequencies between the GWAS variants and all variants in the selection scan were similar (Fig., S3.3). Nevertheless, to control for potential bias, MAFs of variants were split into bins of size 0.025. We calculated the density of our overlapping GWAS and selection variants using a bin size of 0.025 (i.e., what proportion of our observed SNPs had a MAF between 0-0.025, 0.026-0.050, etc.). These densities were used to weight random sampling of 139 selection scan variants (the number of loci identified as GWAS outliers) to simulate the MAF distribution in the GWAS variants. We then calculated how many of these selected variants overlapped with outlier regions. This was repeated 2500 times and our observed values were compared to this null distribution and p-values were calculated (Fig. 4B). Finally, we used Bedtools to identify the closest gene to each outlier SNP using the gene models from Reid et al. [36].

## Results

### Phenotypic variation

There was significant variation in each of the physiological phenotypes measured both within and between populations (Fig. 2). In low salinity, fish must actively regulate their internal homeostasis to prevent ion loss and limit water influx [55]. Fish that fail to acclimate will experience a dilution of body fluid and individuals with greater tolerance will maintain higher internal ion concentrations. For our freshwater challenge, parental populations did not differ significantly in plasma Cl^-^ regulation (all results mean ± standard error; BW-native = 122.0 ± 1.46 mM; FW-native = 126.5 ± 1.64 mM; p = 0.13), but qualitatively the FW-native population maintained higher plasma Cl‐ when facing freshwater challenge than BW-native individuals, consistent with previous research that showed similar but significant differences between these populations [20,3]. Admixed individuals lost Cl^-^ homeostasis to a greater degree than either of the parental populations (admixed = 108.1 ± 0.84 mM; p < 0.001; Fig. 2A), a pattern that is consistent with negative epistasis. Hypoxia tolerance showed a clinal pattern where FW-native individuals tolerated low oxygen levels for the shortest time (30.3 ± 2.81 min), BW-native for the longest time (89.5 ± 2.71 min), and admixed individuals were intermediate (50.0 ± 1.64 min; Fig. 2B). All population comparisons were significant at p < 0.001. MR results differed from hypoxia tolerance; FW-native individuals had high MR (4.7 ± 0.20 μmol O_2_g^-1^hr^-1^), yet admixed (3.3 ± 0.03 μmol O_2_g^-1^hr^-1^) and BW-native populations (3.2 ± 0.07 μmol O_2_g^-1^hr^-1^) were significantly lower (p < 0.001) though not different from each other (p = 0.96). Finally, regressions between the three phenotypes in admixed individuals revealed no significant correlations, suggesting that phenotypic variation in salinity tolerance, hypoxia tolerance, and MR are inherited independently (p>0.56 for all comparisons).

**Figure 2.**
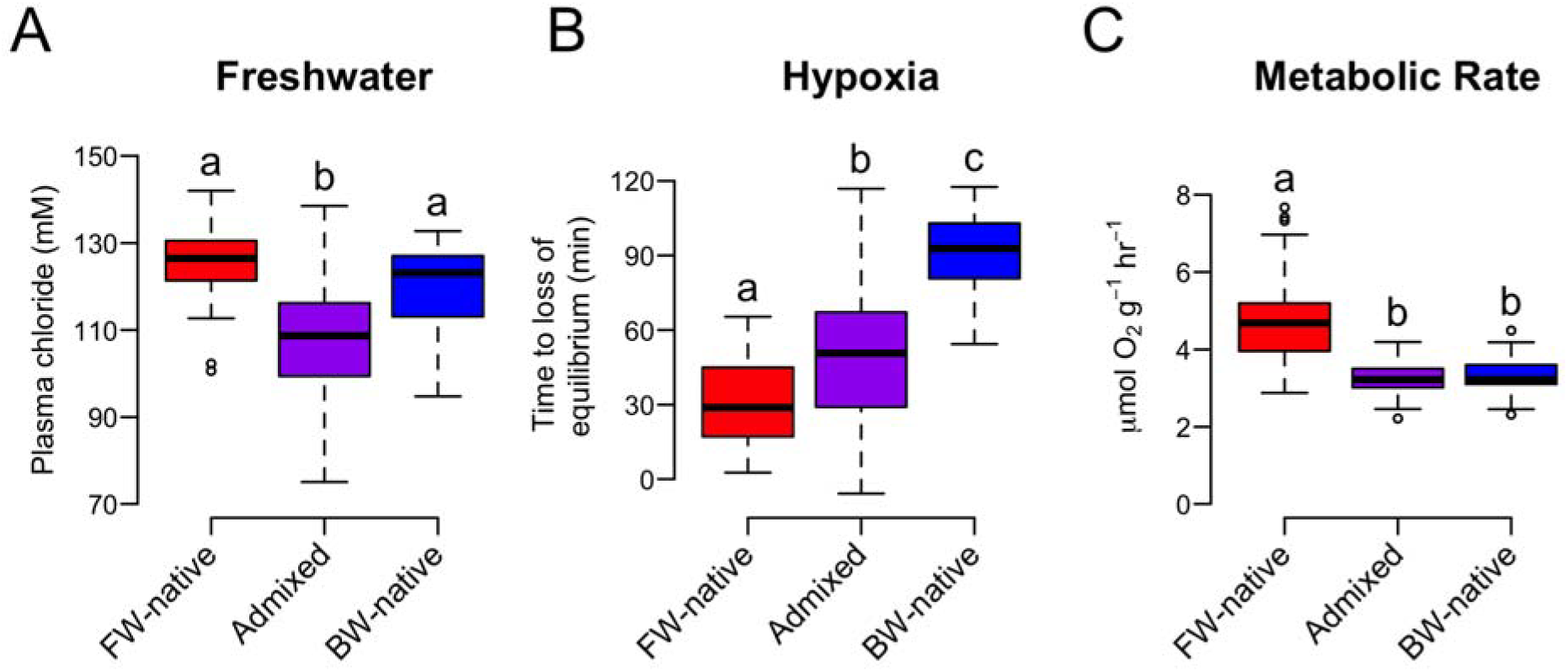
Variation in physiological traits for all populations. Plots depict freshwater tolerance (A), hypoxia tolerance (B), and metabolic rate (C) for the FW-native (red), admixed (purple), and BW-native (blue) populations. Data presented in Tukey boxplots and all values have been mass corrected to a fish size of 4g (for visualization purposes only). Letters above each box indicate statistically significant differences following post hoc tests.

Morphological data suggest that all populations are distinct. PCA separates the admixed individuals from both parental populations along PC1 (46% of the variation) while PC2 distinguishes the BW-native and FW-native populations (17% of the variation) (Fig. S6). The shape changes between populations appear to depend on the relative location of the caudal and anal fin relative to the tail for PC1; there is also a subtle shift in eye location. PC2 suggests similar shifts; however, the ratio of the tail fin itself is also elongated and shortened in FW-native versus BW-native individuals. Permutation tests from the CVA reveal significant differences for Mahalanobis distances for all populations (p < 0.001) but Procrustes distance permutations show only differences between admixed and the two parental populations (p < 0.0001) but no difference between the parental populations (p = 0.065).

### Population genetics

Population genetic results demonstrate that our admixed population is genetically intermediate to both parental populations (Fig 3A, B). Principle component (PC) 1 explains 20% of the variation among individuals and distinguishes populations along the salinity cline. PC2 separates admixed individuals from both parental populations, and explains 7% of the individual variation (Fig 3A). For admixture analysis, genotypes from parental populations clustered into different groups while individuals sampled from the zone of admixture have an average admixture proportion of 0.62 (range 0.52 to 0.70) (Fig. 3B). Genome-wide average F_ST_ values between populations were: FW-native vs. BW-native 0.082, FW-native vs. Admixed 0.035, BW-native vs. Admixed 0.022.

**Figure 3.**
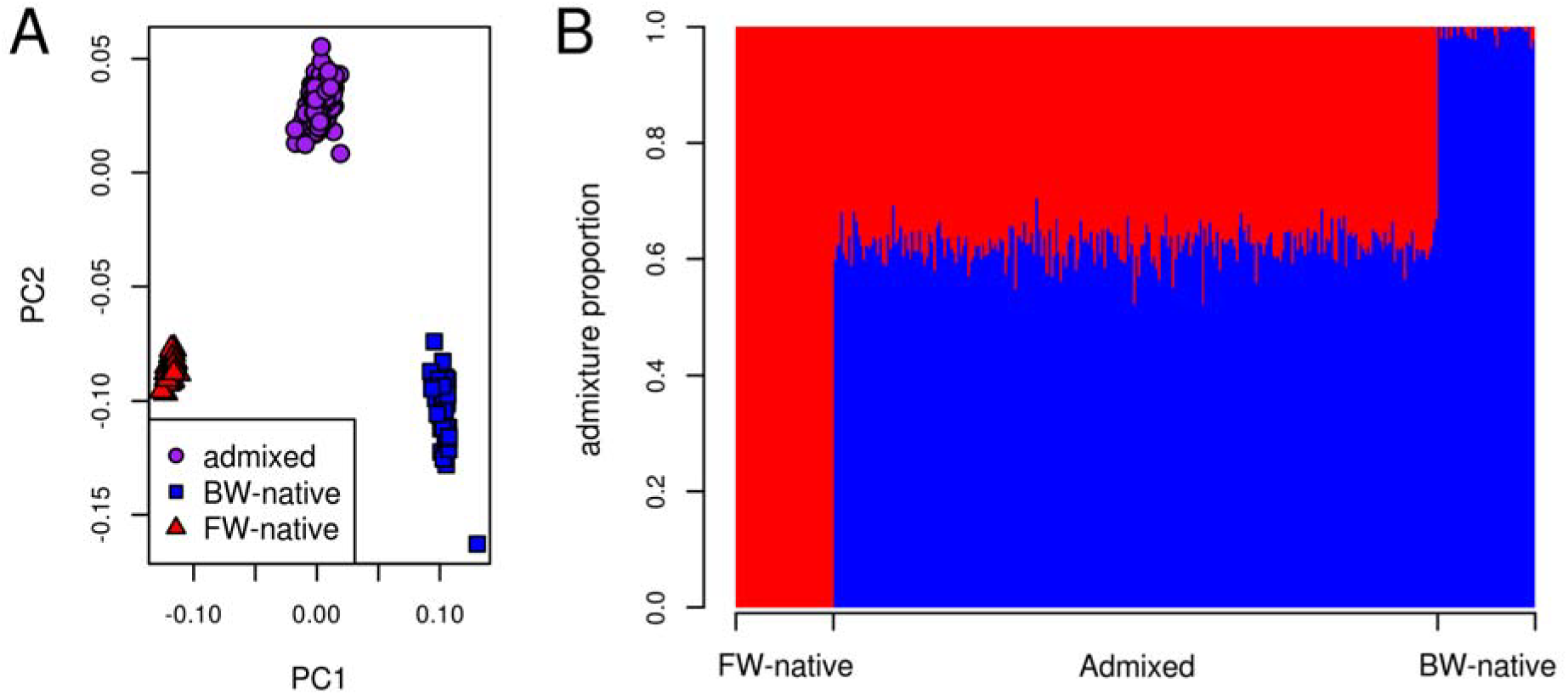
Population structure of the FW-native, admixed, and BW-native populations as determined by PCA (A) and Admixture (B) analyses. (A) Each point represents an individual genotype where shapes and colors distinguish populations. The amount of variation explained by PC1 and PC2 is 20% and 7%, respectively. (B) Inferred ancestry proportions for each individual where the number of ancestral populations (K) is 2. Each bar represents an individual and x-axis location specifies sampling location. Blue and red represent BW and FW ancestry, respectively.

### GWAS

Association results using typical genome-wide Bonferroni-corrected significance cutoffs yielded significantly associated loci for only the MR phenotype, which had 1 SNP passing the threshold (Fig. S2). Hypoxia tolerance had 1 variant passing the LD thinned threshold. The 0.1% most significant GWAS hits produced 139 loci for each phenotype and these were distributed throughout the genome (Fig. S4).

### Selection scans

Selection scans revealed regions of elevated divergence across the genome. We identified 1387 outlier loci when using an LD value of 27 kb. These loci collapsed into 679 outlier windows with an average size of 3.5 kb. 285 of the windows contained more than 1 outlier SNP. The largest outlier window contained 18 outlier variants with a length of 128.9 kb, falling on chromosome 8 from 2.61-2.74 Mb. See figure S1 for distribution of outlier window lengths.

### Overlap between GWAS and selection scans

We identified 16 outlier physiological GWAS variants that fell within genomic regions showing signatures of natural selection between marine and FW populations. Permutation tests show that these top phenotype-associated loci were more likely to show signatures of selection than by chance (Fig. 4B). That is, genomic signatures of selection were enriched for top GWAS loci for each of the physiological traits. Salinity and hypoxia showed 6 overlapping variants between GWAS and selection outlier regions (p = 0.005; p = 0.002). The 4 overlapping loci for MR were also more than expected by chance (p = 0.05). See Table 1 for a complete list of overlapping variants. Of the 16 candidate SNPs, 13 fell in coding regions of genes while the rest were in non-coding regions. Morphological analysis showed only 3 and 2 overlapping variants between GWAS and selection scans, which is no more than is expected by chance (p > 0.05).

**Figure 4.**
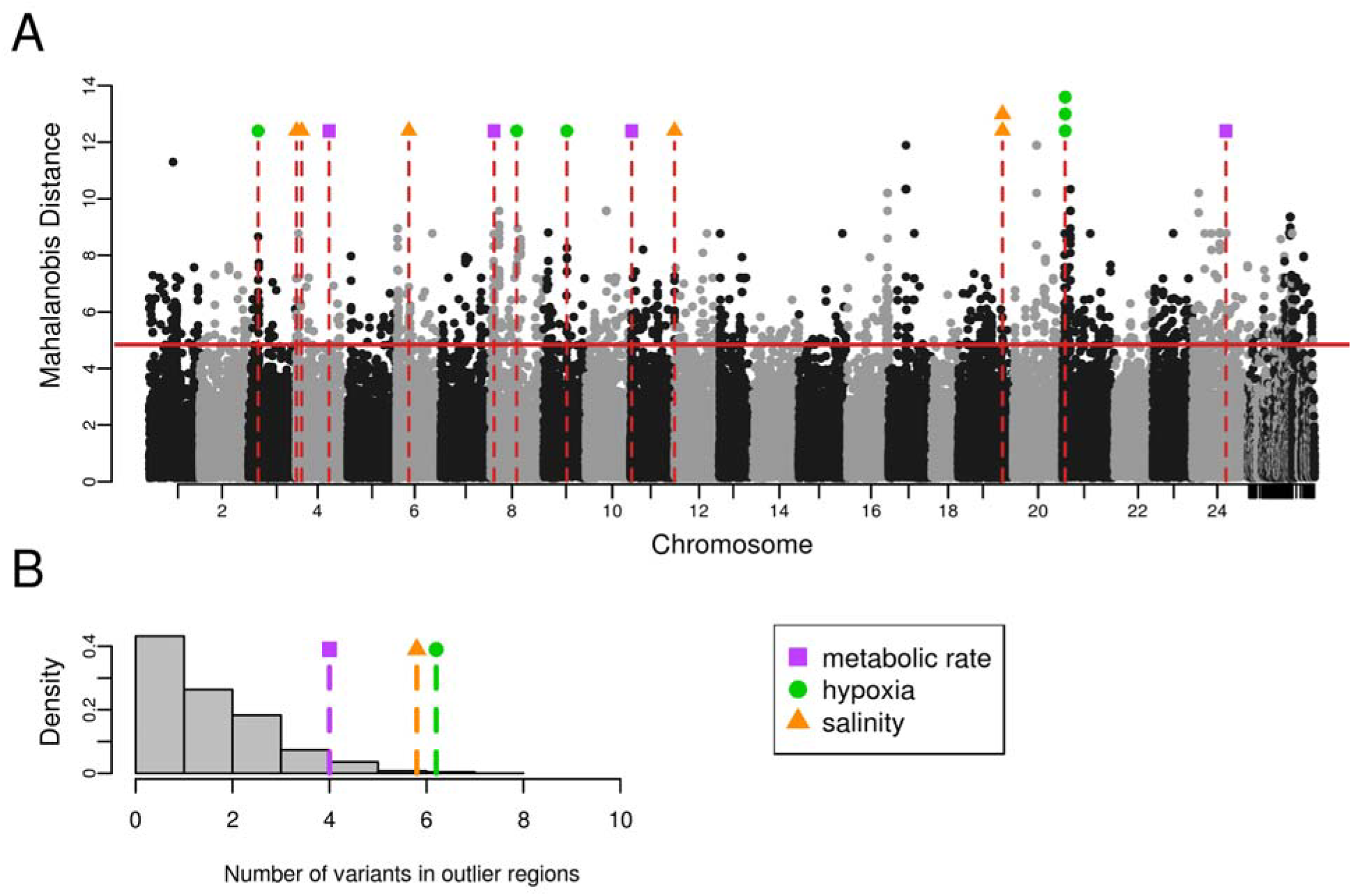
Integration of genome-wide scans for signatures of natural selection and genome-wide association mapping of physiological traits. (A) Signatures of selection as represented by Mahalanobis distance across the genome, where alternating black and gray dots represent genetic variants associated with different chromosomes. Horizontal solid red line is the 99% cut-off for the selection scan. Each black or grey point represents a variant and its location in the genome. Unplaced scaolds are placed on the far right. Vertical dashed red lines signify overlap between loci associated with phenotypes (from GWAS) and loci showing signatures of selection between BW-native and FW-native populations. The color of the symbol above each line specifies the relevant phenotype. Where multiple variants are associated with phenotype and fall within a single region showing a signal of natural selection, points are stacked. (B) Results from 2,500 permutations to test for non-random overlap between phenotype-associated variants with variants showing signatures of natural selection. Grey bars represent the distribution of the number of variants expected to overlap from random permutations. Each shape and dashed line are the number of empirically-discovered overlaps. Note that salinity and hypoxia have been jittered for clarity, but both have 6 overlapping variants. The number of trait-associated variants that also show signatures of natural selection are greater than expected by chance, for each of salinity, hypoxia, and MR (p=0.005; p=0.002; p=0.05).

**Table 1.**
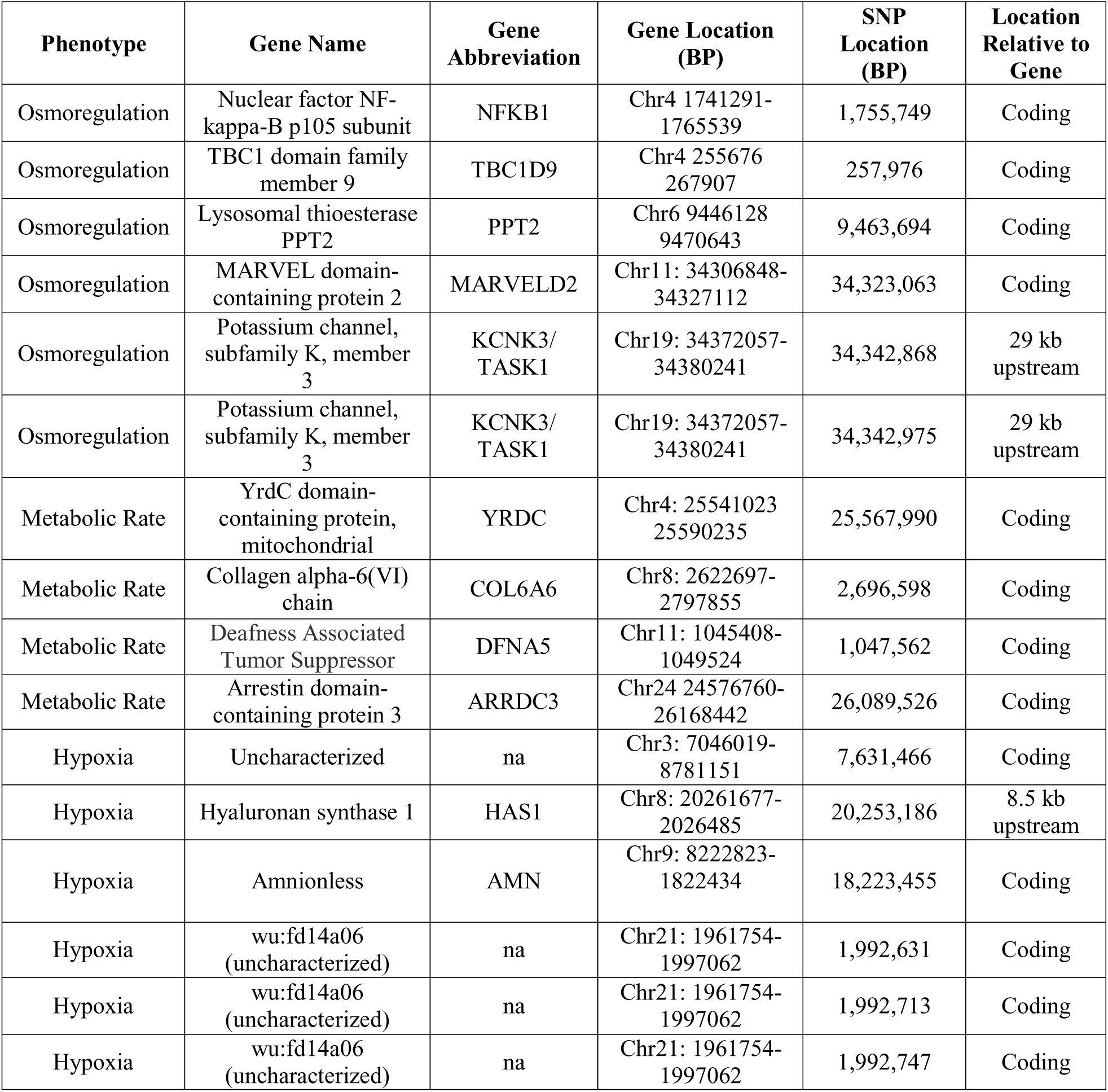
Candidate SNPs underlying the adaptive phenotypes in *F. heteroclitus*. All variants listed are found in outlier selection scan regions and a top association hit. The phenotype column specifies the phenotype used for the association mapping.

It should be noted that the method for LD estimation slightly influences our results. Using the 27 kb LD cutoff, our variants are informative of ∼73% of the genome. The more conservative 5 kb cutoff provides information for ∼25% of the genome and the outlier windows collapse into 839 windows with an average of 0.2 kb and a max of 4.8 kb. However, this only slightly alters our main findings. Hypoxia tolerance, salinity tolerance, and MR drop to 4, 4, and 2 overlapping association and selection variants (p = 0.05, p = 0.04, p = 0.31). The genes that no longer overlap for MR are ARRDC3 and DFNA5. Hypoxia outliers no longer contain the uncharacterized protein and AMN while salinity no longer contains TBC1D9 and MARVELD2.

## Discussion

When studying adaptation to a complex environment, phenotypes that are the targets of natural selection may be many and complex. For inferring the genetic basis of such adaptations, phenotype agnostic approaches such as genome scans provide a path forward, but interpreting the phenotypic relevance of the implicated genes can be difficult. Conversely, approaches such as GWAS focus specifically on the variants underlying phenotypes of interest, but provide no information concerning the adaptive significance of these variants. We attempted to confront and ameliorate the limitations of these two approaches by integrating them; an idea that has only recently begun to gain traction [53,56-59]. Our approach is unique in that we leverage LD generated by natural admixture between populations to identify genotype-phenotype associations (see also Brelsford *et al.*, 2017). We first located potential adaptive regions of the genome using scans for selection then used GWAS to reveal genomic regions that are associated with multiple potentially adaptive phenotypes. By identifying the regions where the two approaches overlap, we isolate the genomic regions under selection while simultaneously inferring physiological function of these regions. At the same time, we provide additional evidence that natural selection is governing the divergence of these physiological phenotypes between marine and freshwater environments. This study extends our understanding of the genotypes and phenotypes that are important for adaptation to alternate osmotic environments, which has heretofore been primarily focused on morphological traits. We interrogate multiple, complex physiological phenotypes that are known to diverge between freshwater and marine environments. This focus enables us to identify mechanisms of adaptive divergence that have likely been important in enabling speciation events across osmotic boundaries in fishes.

We find evidence that salinity tolerance, MR, and hypoxia tolerance are diverging through adaptive processes. There is significant enrichment in overlap between top-ranked GWAS loci for each phenotype and top-ranked selection scan regions; this suggests that the regions of the genome influencing key phenotypes that are known to vary between osmotic niches are disproportionally divergent between the FW-native and BW-native populations (Fig. 4). This finding, coupled with the physiological divergence between populations (Fig. 2), is consistent with adaptive divergence in salinity tolerance, MR, and hypoxia tolerance; we conclude that divergence in these three physiological traits is important for enabling evolutionary transitions from marine to freshwater environments.

In stickleback and alewife, body shape diverges between marine and freshwater populations [27,60] [27] and Baltic Sea herring morphology shifts due to salinity adaptation [61]. As such, we hypothesized that body shape may differ between *F. heteroclitus* populations in different salinities. We identify subtle shifts in body shape between environments, though this phenotype does not appear to be under selection. While divergence in morphological characteristics is adaptive in some fish systems, this does not seem to be the case for BW-native and FW-native populations of *F. heteroclitus* (supplemental information). There are likely other selective pressures that are important for freshwater adaptation including pathogens, diet, microbiome, predator regime, etc., and future work should begin to integrate these additional complexities.

Physiological and genomic results suggest that there is limited overlap in the genetic underpinnings of the three physiological traits. Co-variance between traits could be due to LD between underlying genes, epistatic interactions between genes, or by pleiotropy, where one gene influences multiple phenotypes [62,63]. We find no correlations between the phenotypes in admixed individuals. Similarly, there is no overlap among adaptive candidate genes between the three phenotypes. As such, the lack of correlations between the phenotypes in admixed individuals suggest that there is minimal shared genetic basis influencing these traits. A shared genetic basis between traits can limit or accelerate the efficiency of natural selection and adaptation, depending on the direction of correlations (positive or negative). The modular nature of these phenotypes implies that each trait can be altered independently, thereby promoting the evolvability of a diversity of complex physiological phenotypes [64,65].

### >Genetic basis of divergence in osmoregulatory physiology

The variation between populations in the ability to regulate plasma chloride suggests that, while the parental populations are adapted to their native salinity, admixed individuals have reduced osmoregulatory capabilities relative to the parents (Fig. 2). This pattern may represent hybrid breakdown due to negative epistasis [66], which is indicative of Bateson-Dobzhansky-Muller incompatibilities [67-69]. Sets of parental alleles that typically do not co-occur due to purging through natural selection may be brought together in hybrids. Because evolutionarily derived alleles tend to be those responsible for incompatibilities [70], adaptation to freshwater may be resulting in the reduction of performance seen in admixed individuals. This process would impede gene flow across the hybrid zone and enable adaptive divergence between the two parental populations.

In fish, gills function as the primary site of osmoregulation. Upon transitioning from high to very low salinity, killifish reversibly remodel their gill morphology as part of a strategy to maintain osmotic homeostasis. This process involves regulating cell volume, replacing or remodeling the morphology of ionocytes, expressing appropriate ion transport proteins and inserting them in the appropriate surface (apical or basolateral) of ionocytes, and modulating junctions between epithelial cells to prevent or allow ion loss [71,72]. Previous work in *F. heteroclitus* has established that the ability to transition between alternate gill morphologies is essential during acclimation to different osmotic conditions [23,30,73]. More specifically, previous comparative physiology and transcriptomics research led to a prediction that adaptive divergence between marine and FW killifish is linked to divergence in both cell junction regulation and ion transport [20,3]. Consistent with this *a priori* prediction, here we identify five genes involved in these processes. These genes (Table 1) are associated with variation in osmoregulatory physiology among individuals, and show signatures of selection between BW-native and FW-native populations, and are therefore likely contributing to adaptive osmoregulatory divergence.

TASK1 is an outwardly rectifying K^+^ channel and may be involved in cell volume regulation. This gene plays a role in shark osmoregulation; it establishes the driving force for apical Cl^-^ excretion via CFTR [74]. CFTR is an important ion channel in killifish at high salinity [75,76], but our experiment focused on acclimation to low salinity, and it is unlikely that an outwardly rectifying K^+^ channel is functioning in the same manner here. TASK2 channels have been found to influence cell volume regulation, particularly in response to hypotonic stress and cell swelling [77]. Regulatory volume decrease (RVD) involves the activation of Cl^-^ and K^+^ channels [78]. It is possible that in *F. heteroclitus* orthologs to TASK1 facilitate RVD and variation in structure of regulation of this gene results in differential osmoregulatory capabilities. This hypothesis merits further investigation.

An intriguing connection exists between adaptive candidate genes NFKB1, TBC1D9 and PPT2. NFKB1 has been implicated as a hub gene regulating the transcriptional response to low salinity acclimation [23,30]. This gene may be involved in the apoptotic processes driving gill remodeling during acclimation. Similarly, TBC1D9 and PPT2 mediate the transport and localization of proteins. Functionally, these three genes may be interacting to enable cellular remodeling to remove and install the appropriate proteins necessary for maintaining osmoregulatory homeostasis in dilute fresh water.

Tight junctions form at intercellular junctions and work to regulate the flow of solutes through the paracellular pathway [79]. Tight junction proteins are well-known components of the osmoregulatory apparatus in fish [71], and have been identified as regulators of variation in osmotic tolerance in stickleback [31], killifish [80], alewives [81], silverside [82], and sea bass [83], among others. At low salinity, tight junctions prevent the loss of Cl^-^ while leaky junctions at high salinity enable the excretion of Na. Tri-cellular junction proteins, which occur where any three cells join, influence paracellular permeability in fish gills [84,85]. MARVELD2 (also known as tricellulin – a tri-cellular tight junction protein) was identified as a target of selection and GWAS outlier, suggesting that tri-cellular tight junctions are likely important in adaptive divergence in osmoregulatory physiology between FW-native and BW-native populations. This adds mechanistic insight into previous research identifying an evolutionary trade-off between BW-native and FW-native populations in the physiological ability to regulate tight versus leaky junctions during osmotic acclimation [23].

We measured the ability to maintain plasma chloride homeostasis as a representation of the phenotypic variation in osmoregulatory abilities among individuals. However, other components of ionoregulatory physiology may be subject to adaptive divergence between marine and freshwater populations. Indeed, many genes show signatures of selection between parental populations that are consistent with this expectation. For example, outlier loci from selection scans are enriched for GO terms *tight junction*, *regulation of intracellular pH*, and *bicarbonate transport*. These include a number of genes that are core components of fish gill ionocytes, including multiple sodium bicarbonate cotransporters (NBC), sodium potassium ATPase (NKA), sodium hydrogen exchanger (NHE), carbonic anhydrase (CA), anion exchange protein (SLC4A3), rhesus glycoprotein (RHCG), and aquaporin (Table S5). This is consistent with an evolved sodium uptake metabolon in the FW-native population, where apical sodium uptake is through NHE and CA, and basolateral sodium export is through NBC and NKA (Figure 5), as reported in rainbow trout [86]. Rhesus glycoproteins interact with the same molecular machinery to pair ammonia excretion with sodium uptake in larval freshwater fish [87]. Outliers also include multiple tight junction proteins (claudins and occludin) that may be involved in adaptive changes in paracellular ion transport pathways. We hypothesize that derived mechanisms of sodium uptake, perhaps paired with ammonia excretion, are an important component of adaptive divergence between marine and freshwater populations of killifish.

**Figure 5.**
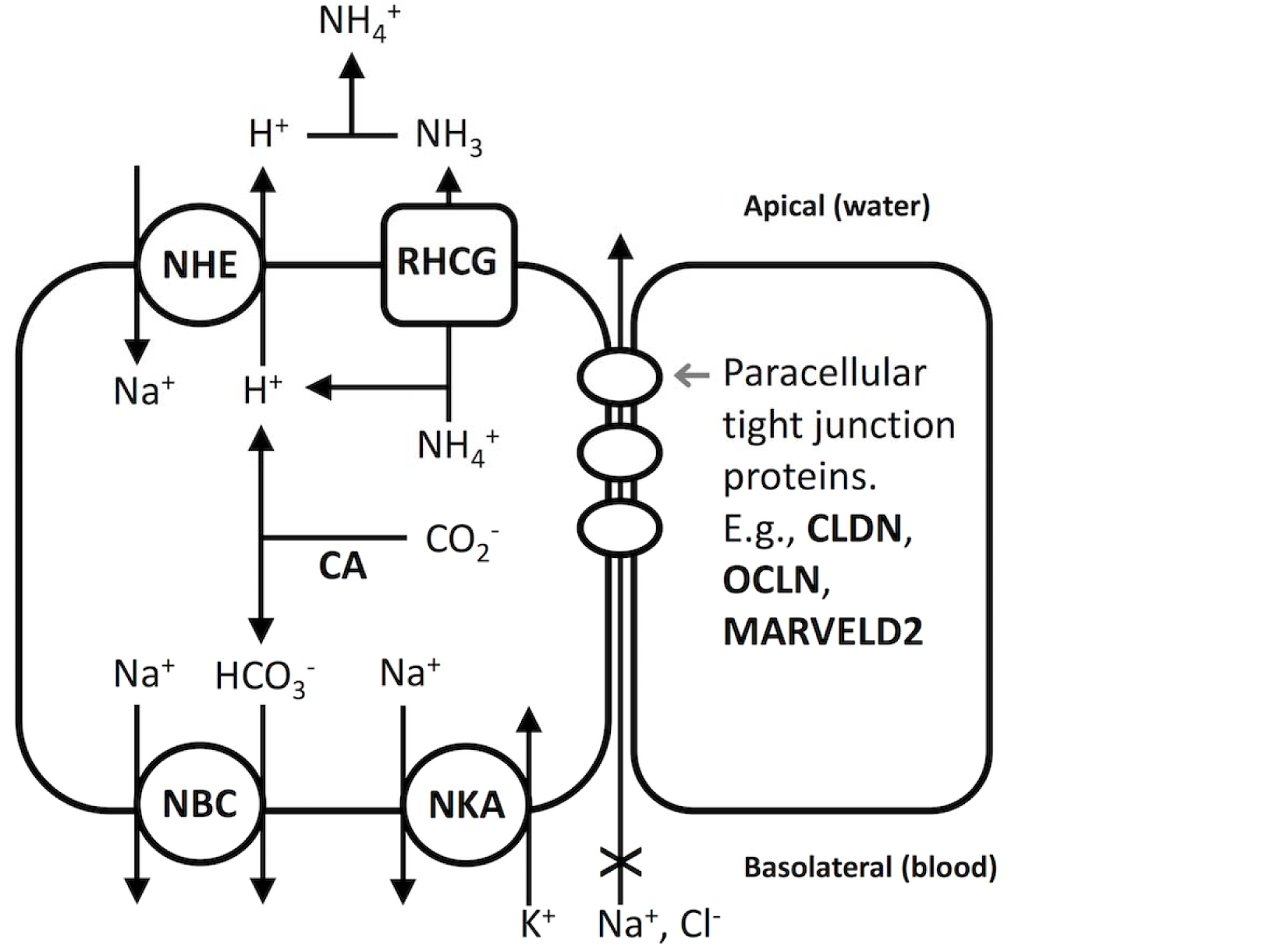
Hypothesized model of adaptive sodium uptake metabolon in freshwater fish. Genes (bold black) that show signatures of selection in genome scans and that contribute to ionoregulation. The model on the left is a fish ionocyte, where proteins constitute a metabolic unit that regulates sodium uptake in fresh water. NHE and CA facilitate sodium uptake from water, and NBC and NKA facilitate export of sodium into the blood. RHCG may also work with NHE to facilitate both ammonia excretion and sodium uptake. The ionocyte and an adjacent cell (right) are joined with paracellular tight junction proteins that regulate paracellular permeability.

### Genetic basis of divergence in metabolic rate

Population divergence in MR, coupled with overlap between GWAS loci and loci under selection, suggests that metabolic rate is diverging by natural selection in different osmotic environments. We identify genetic variation in four genes that contribute to adaptive divergence in metabolic rate between populations. Yet, the adaptive significance of metabolic rate divergence between osmotic environments is unclear. For example, this phenotype does not seem to correspond to swim performance [22]. However, our results suggest that further study of the adaptive relevance of alternate metabolic rate physiologies between marine and FW environments is merited.

We identify four genes that are associated with inter-individual variation in MR, and that are evolving by natural selection between marine and FW environments (Table 1). Knockout studies in mice have shown that ARRDC3 and COL6A6 regulate MR. Both genes have been directly linked to regulating oxygen consumption and energy use [88-90]. DFNA5 is a gene involved in hearing loss in humans. However, knockout mice for this gene show a down-regulation of genes involved in energy metabolism, suggesting a role in regulating metabolic rate [91]. Though there has been relatively little work linking candidate YRDC to metabolism, since it controls mitochondrial tRNA synthesis [92] it is plausible that it may influence energy production. Healy *et al*., [93] showed that YRDC expression is increased in cold temperatures in coastal populations of *F. heteroclitus*. Increased metabolic activity at low temperatures is necessary to ensure sufficient energy available for cellular functions [94] and the up-regulation of YRDC at low temperature may indicate a link with MR.

### Genetic basis of divergence in hypoxia

Hypoxic environments occur regularly in marsh systems, primarily due to isolation of pools (i.e., during tidal cycles) and nocturnal respiration [95-97]. However, the oxygen availability in fresh versus salt tidal wetland differs where low salinity habitats have higher oxygen concentrations than those of higher salinity [12]. Therefore, one might predict that BW-native and FW-native populations regularly experience environments with different oxygen availability, and have diverged in their relative ability to acclimate to hypoxia challenge. Population differences are consistent this expectation where FW-native individuals harbor lower hypoxia tolerance than the BW-native fish (Fig. 2).

We identify genetic variation in six genes that contribute to adaptive divergence in hypoxia tolerance between populations. Of these, four are uncharacterized, making biological interpretation difficult. AMN is a transmembrane protein that is important during development, but has no clear connection to hypoxia [98]. However, we identify a variant 8.5 kb upstream of HAS1 as a promising candidate. HAS1 synthesizes cellular hyaluronan, which is a polysaccharide that is one of the main components of the extracellular matrix [99]. Hyaluronan has been directly implicated as a negative regulator of hypoxia-inducible factor-1alpha (HIF‐ 1alpha) under hypoxic conditions [100]. HIF genes are transcriptionally responsive to hypoxia in *F. heteroclitus* [101,102]. If variation in HAS1 influences the production of hyaluronan, this could lead to a direct effect on hypoxia tolerance.

## Conclusion

Selection scans are a powerful method to identify putatively adaptive loci between populations diverging across different environments. However, determining the phenotypic effects of these loci is dependent on inferring function from annotated genes. While this approach is a useful first step, it is limited by the fact that many genes have species-specific, ambiguous, or unknown functions thereby reducing our ability to link these genes to phenotypes. Making these links becomes even more difficult as multiple, polygenic, phenotypes are involved in the process of adaptation, a characteristic that is likely representative of adaptation to many environments [10]. To overcome these issues, it is necessary to understand the identity and nature of specific phenotypes that are important in adaptation. We addressed this by integrating GWAS and selection scans. This approach was useful for two reasons. First, we were able to identify genetic variation that is both under selection and that explains phenotypic variation. Second, we can begin to disentangle traits that are diverging though adaptive processes in contrast to neutral drift. If phenotypic variation is underpinned with genetic variation that is not under selection, it is likely that neutral rather than adaptive processes are the main drivers underlying population differentiation.

We have demonstrated the strengths of this integrative approach by identifying the genetic variation that is important for adaptation to a freshwater environment. Indeed, nearly all of our identified genes are promising candidates. In particular, the implication of tricellulin as a gene that contributes to adaptive divergence in osmoregulatory physiology is consistent with a priori hypotheses generated by our previous research findings. Focusing on multiple traits allowed us to gain a more integrative understanding of how fish evolve upon radiating into fresh water. Together, these data identify the genetic basis of adaptation to a freshwater environment and demonstrate a promising approach with which to identify the mechanisms important in adaptive divergence between niches.

## Acknowledgments

We thank Michelle Tse and Ruth Hwang for assistance with DNA extractions. Michael Miller and Sean O’Rourke provided training and assistance for RAD library preparation and analysis. David Metzger, Dillon Chung and Kyle Crowther provided assistance with fish collection.

## Data Availability

Sequence data are archived at the National Center for Biotechnology Information (NCBI SRA; BioProject PRJNA428529)

## Author Contributions

Conceived and designed the experiments: RSB TMH PMS AW. Performed the experiments: RSB TMH HJB MVL. Analyzed the data: RSB and TMH. Wrote the paper: RSB TMH PMS AW.

## Supplemental information

**Table S1. Selection scan outlier regions**. Table includes location, closest annotated Fundulus heteroclitus gene annotation, and corresponding UniProt and NCBI annotations.

**Table S2. Salinity tolerance GWAS variants**. Table includes location, closest annotated Fundulus heteroclitus gene annotation, and corresponding UniProt and NCBI annotations.

**Table S3. Metabolic rate GWAS variants**. Table includes location, closest annotated Fundulus heteroclitus gene annotation, and corresponding UniProt and NCBI annotations.

**Table S4. Hypoxia tolerance GWAS variants**. Table includes location, closest annotated
Fundulus heteroclitus gene annotation, and corresponding UniProt and NCBI annotations.

**Table S5. Outlier genes involved in ionoregulation.**

**Figure S1.**
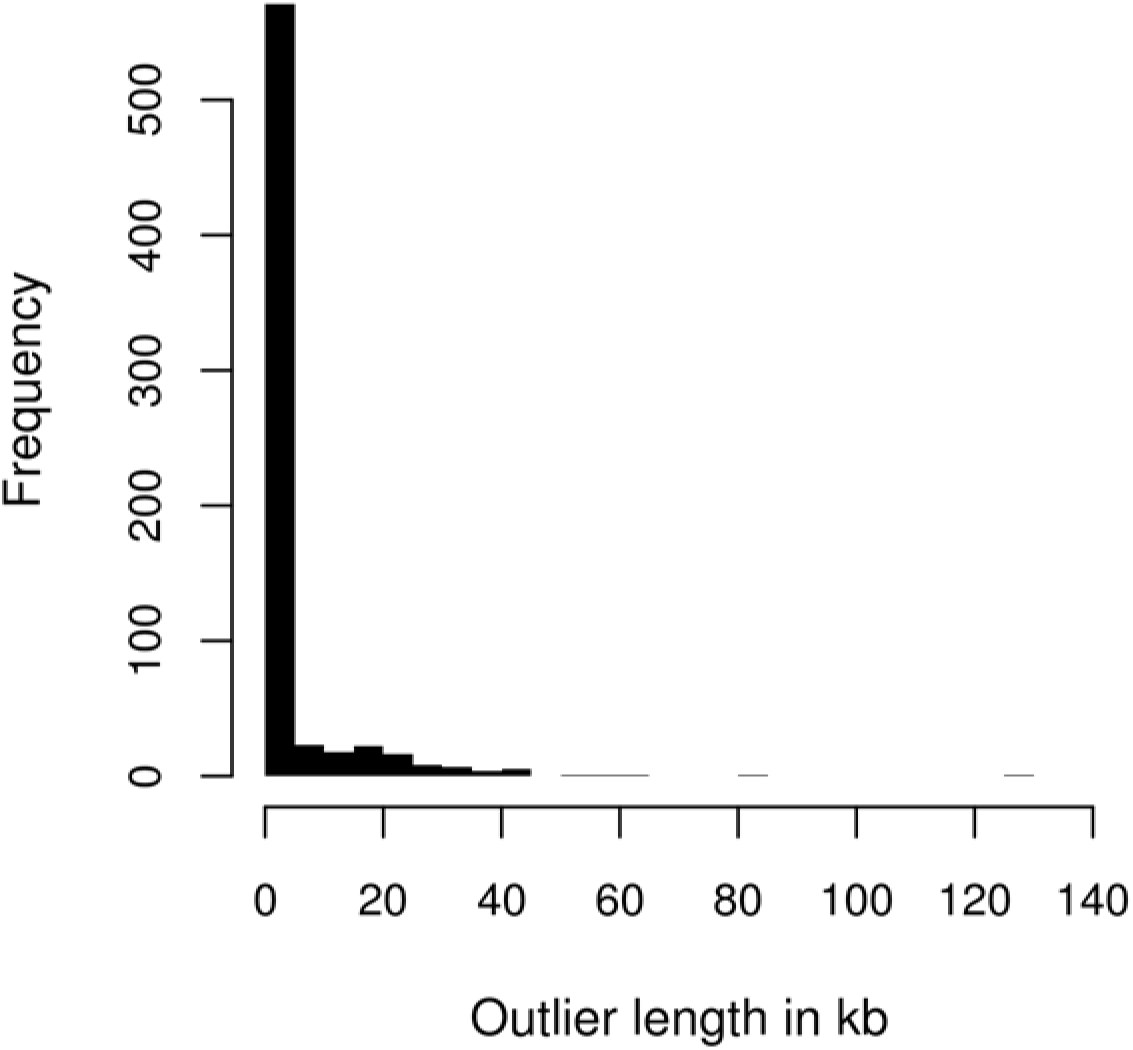
Distribution of the length of outlier regions from the selection scan.

**Figure S2.**
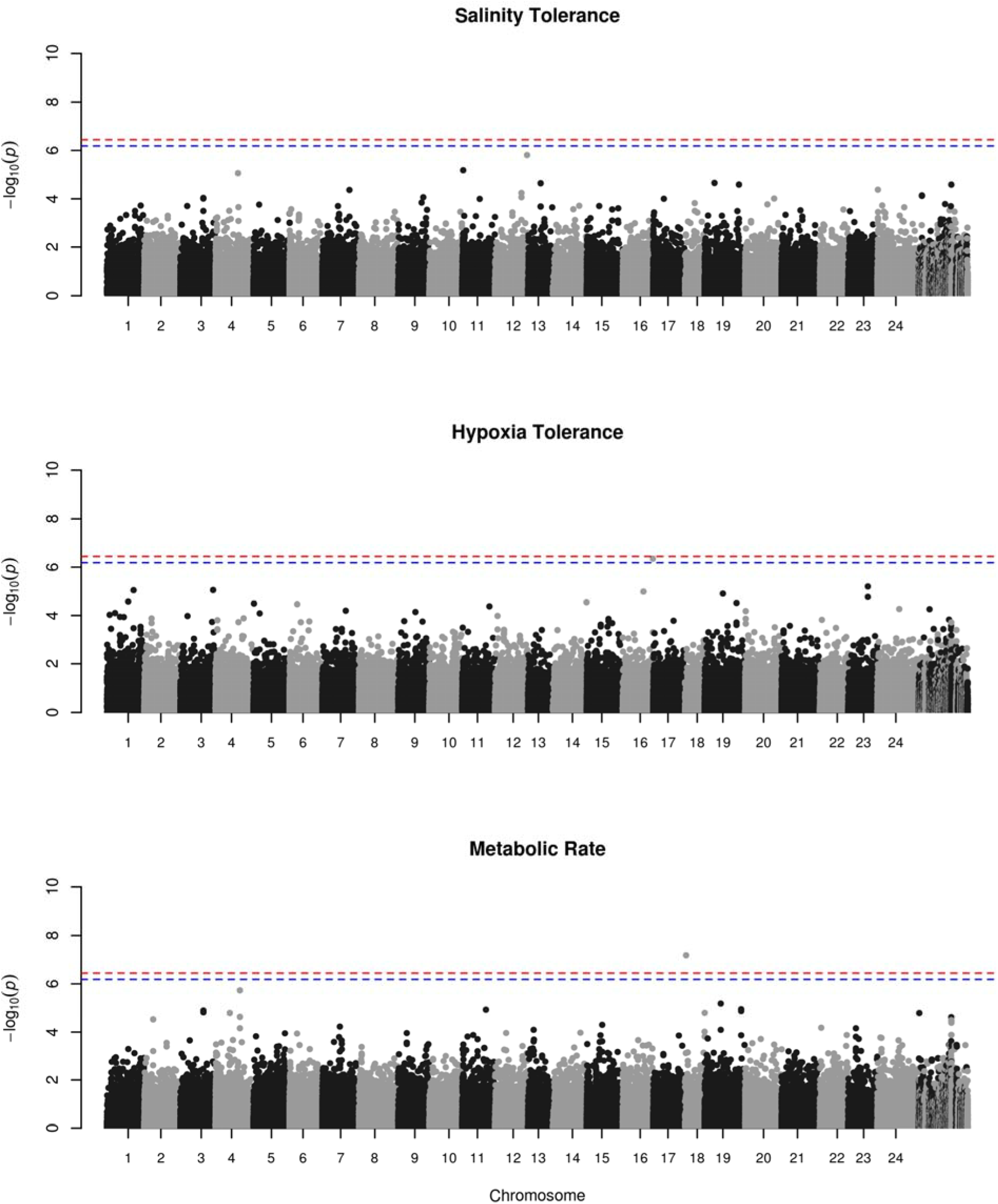
Manhattan plot of association mapping for each physiological phenotype. Dashed horizontal lines indicate Bonferroni corrected significance cutoffs. Red line is the cutoff using the total number of variants while the blue line is a more liberal threshold using the number of LD thinned variants.

**Figure S3.**
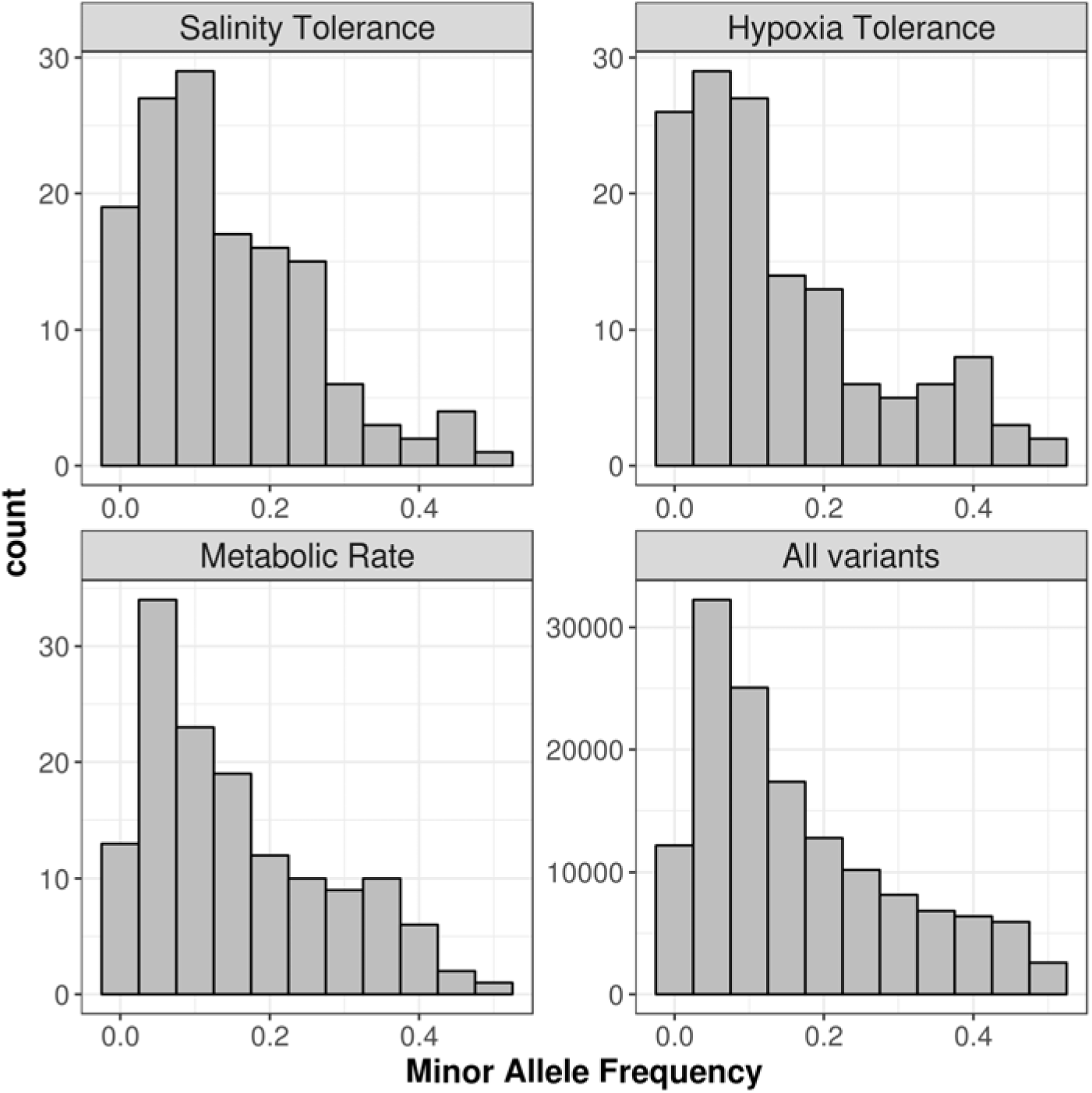
Distribution of minor allele frequencies (MAF) calculated from parental populations only. “All variants” are all SNPs included in the selection scan. The other panels include MAF of the outlier variants from each association mapping phenotype.

**Figure S4.**
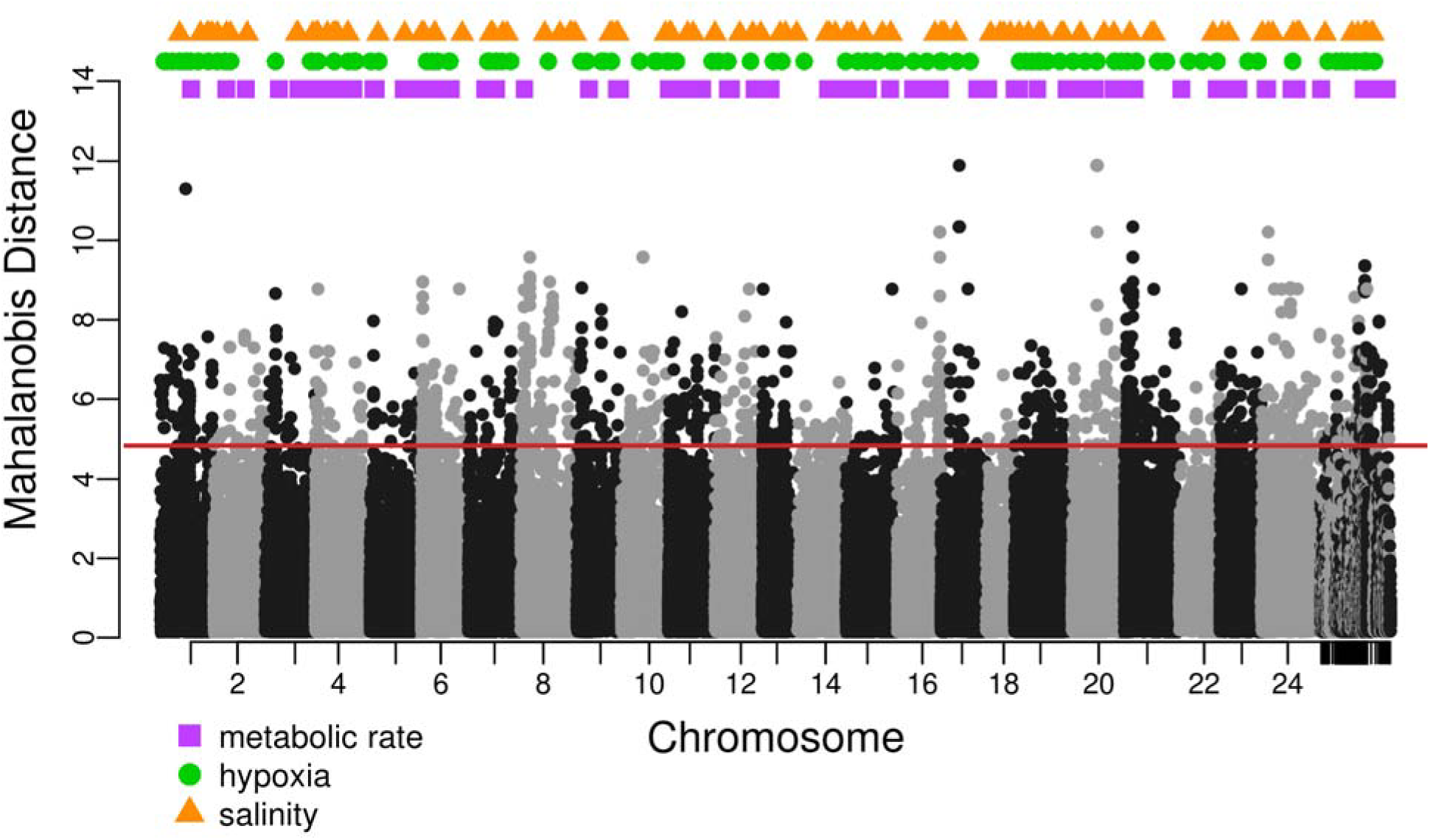
Location of phenotype-associated markers across the genome (top) superimposed on selection scan data. Horizontal solid red line is the 99% cut-off for the selection scan. Each black or grey point represents a variant and its genome position where the y axis position is Mahalanobis distance. Unplaced scaffolds are placed on the far right. Shape and color of the point above the plot specify the association phenotype.

**Figure S5.**
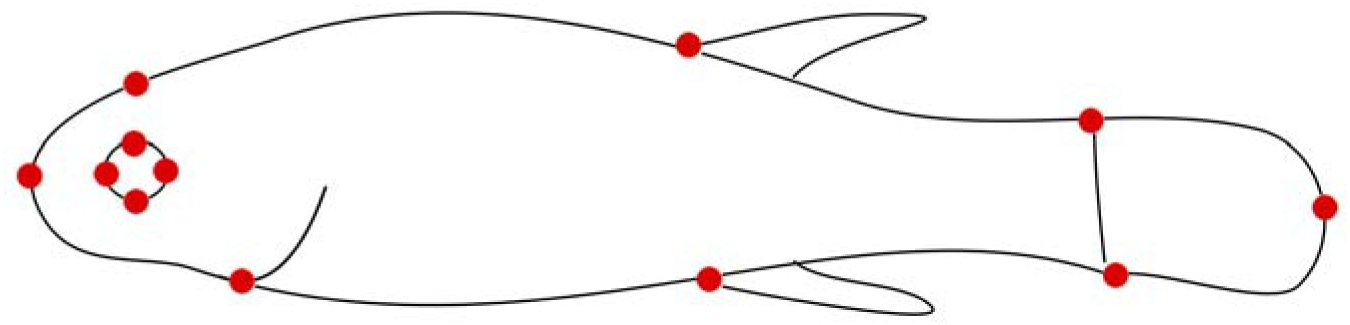
The 12 landmark positions for geometric morphometric analysis.

**Figure S6.**
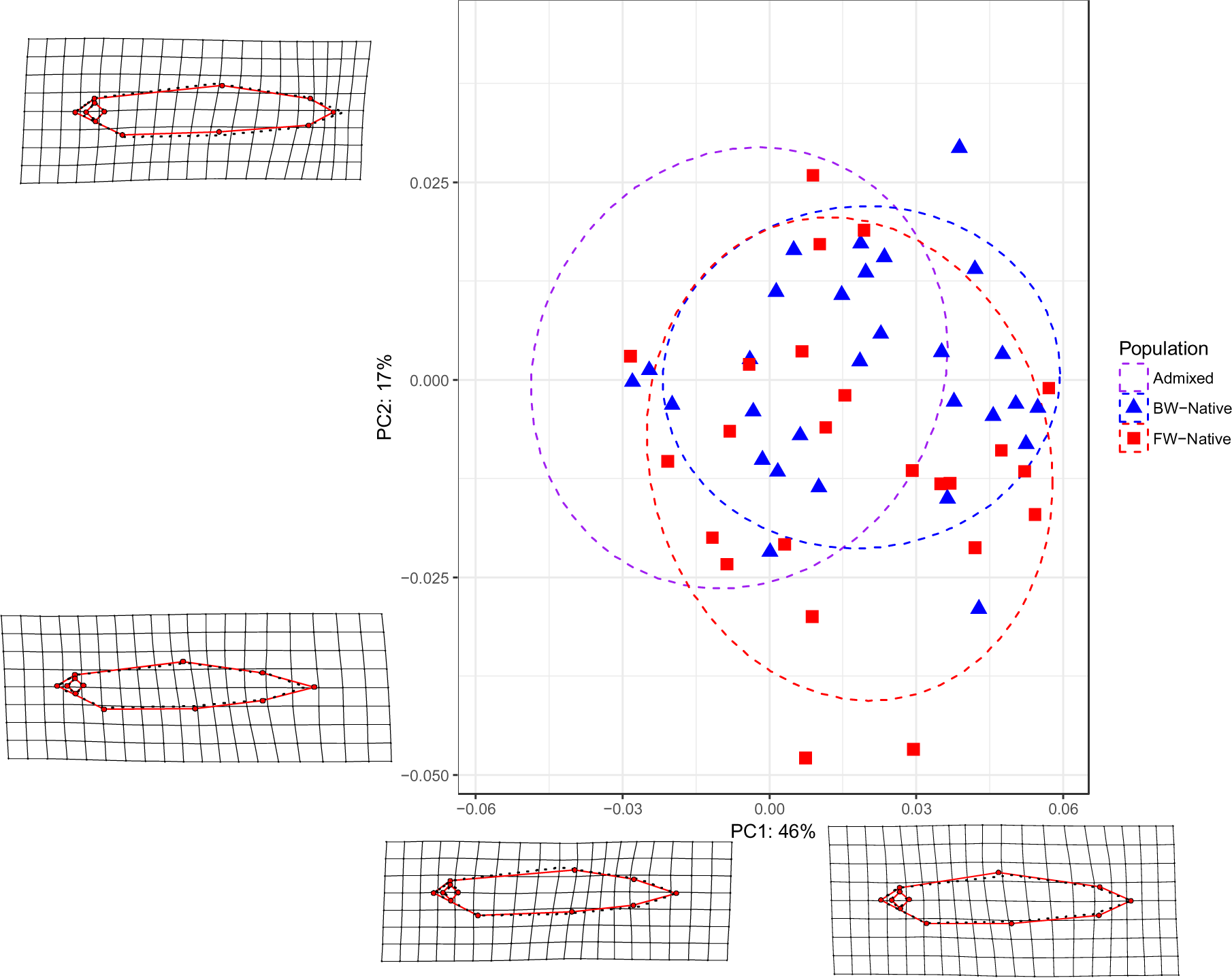
Principal component analysis of body shape. Shape and color indicate population. Dashed ellipses represent 75% confidence intervals, for visualization purposes only. Grid and outline demonstrate shifts in phenotype along each PC axis. Dashed black lines are the average shape while solid red lines show the phenotypic extreme along each axis.

**Figure S7.**
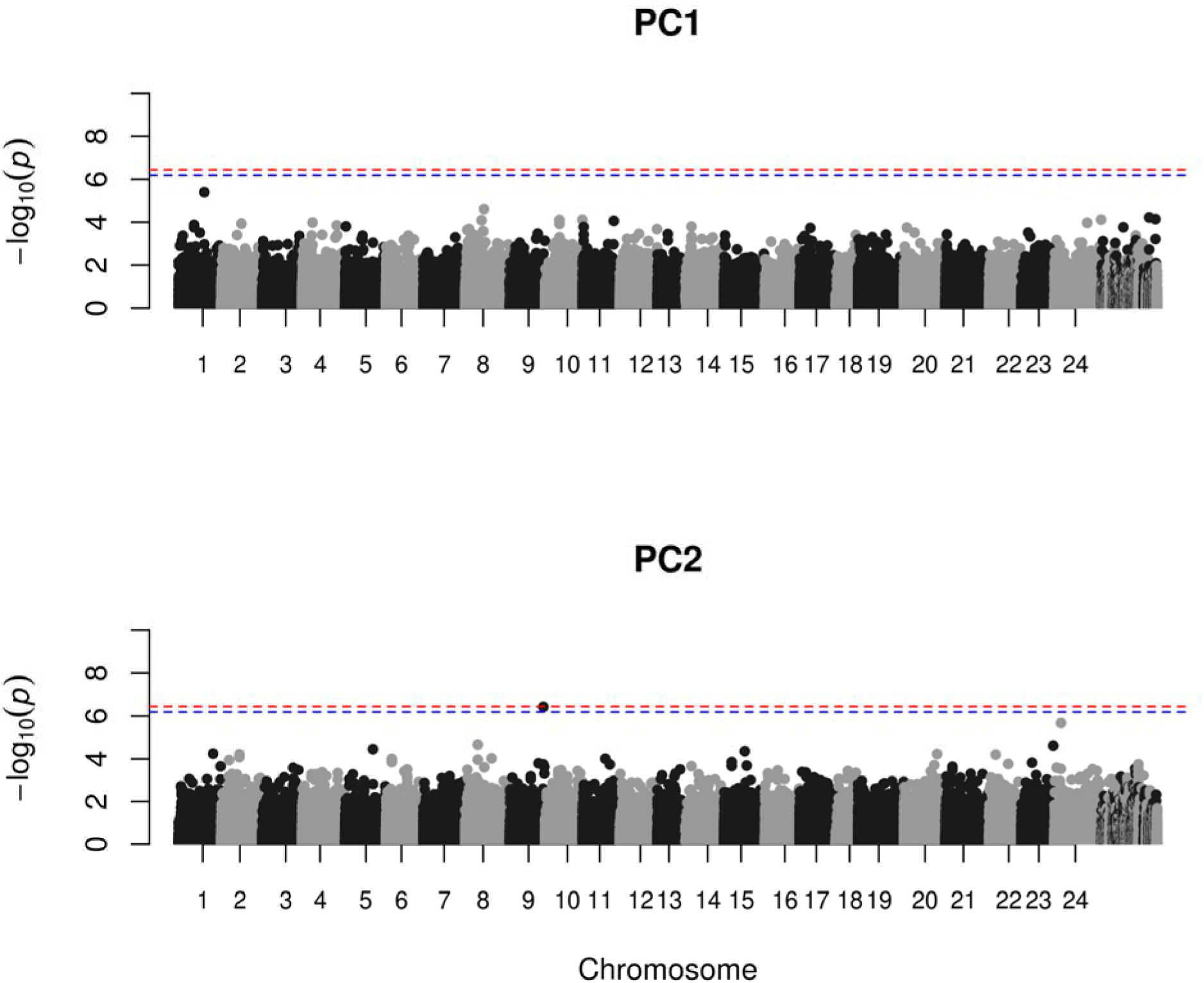
Manhattan plot of association mapping for each morphology principal component. Dashed horizontal lines indicate Bonferroni corrected significance cutoffs. Red line is the cutoff using the total number of variants while the blue line is a more liberal threshold using the number of LD thinned variants.

